# Unlocking a high bacterial diversity in the coralloid root microbiome from the cycad genus *Dioon*

**DOI:** 10.1101/381798

**Authors:** Pablo de Jesús Suárez-Moo, Andrew P. Vovides, M. Patrick Griffith, Francisco Barona-Gómez, Angélica Cibrián-Jaramillo

## Abstract

Cycads are among the few plants that have developed specialized roots to host nitrogen-fixing bacteria. We describe the bacterial diversity of the coralloid roots from seven *Dioon* species and their surrounding rhizosphere and soil. Using 16S rRNA gene amplicon sequencing, we found that all coralloid roots are inhabited by a broad diversity of bacterial groups, including cyanobacteria and Rhizobiales among the most abundant groups. The diversity and composition of the endophytes are similar in the six Mexican species of *Dioon* that we evaluated, suggesting a recent divergence of *Dioon* populations and/or similar plant-driven restrictions in maintaining the coralloid root microbiome. Botanical garden samples and natural populations have a similar taxonomic composition, although the beta diversity differed between these populations. The rhizosphere surrounding the coralloid root serves as a reservoir and source of mostly diazotroph and plant growth-promoting groups that colonize the coralloid endosphere. In the case of cyanobacteria, the endosphere is enriched with *Nostoc* spp and *Calothrix* spp that are closely related to previously reported symbiont genera in cycads and other early divergent plants. The data reported here provide an in-depth taxonomic characterization of the bacterial community associated with coralloid root microbiome. The functional aspects of the endophytes, their biological interactions, and their evolutionary history are the next research step in this recently discovered diversity within the cycad coralloid root microbiome.

## Introduction

Specialized root modifications that contain endophytic microbes are rare in plant evolution, present only in legumes in angiosperms [1] and in cycads in the gymnosperms [2]. Cycads have specialized apogeotropic roots of small coral-like shapes, termed coralloid roots, which contain nitrogen-fixing cyanobacteria. The formation of a coralloid develops from the secondary roots through morphological changes that include an increase in lenticel cells [3], considered the main mode of cyanobacterial entry [3,4]. Coralloid masses are formed involving somatic reduction before the dichotomous branching of the roots. The reduced cells make up part of a ring of differentiated cortical cells lying beneath the epidermis [5]. It is possible to see this ‘cyanobacterial ring’, also known as ‘algal ring’, inside which endophytes are located, even with the naked eye.

Given that all extant cycads have the capacity to form coralloid roots, they were most likely present in the earliest cycad lineages, at least 250 MYA [6]. Compared to the more recent legume nodules of approximately 65 MYA [7], the cycad lineage has a deep evolutionary history of hosting bacterial endophytes in their coralloid roots. Previous investigations of the coralloid roots that trace back as far as the 19th century report bacterial groups belonging to rhizobia [8], *Pseudomonas radicicola* and *Azotobacter* strains [4,9]. Most studies however, have been developed using cyanobacteria-specific markers: morphological and biochemical characters [10,11]; tRNA LEU intron [12,13]; Short Tandemly Repeated Repetitive Sequences [14]; and 16S rRNA direct from the coralloid roots [15], or isolated in BG11 and BG11o media [11,16]. These studies report a simple bacterial community composed of a few or many strains of *Nostoc* cyanobacteria, with one or several strains per coralloid root [12–14]; an overall lack of a geographic structure of the endophyte cyanobacteria in cycad host species from Asia (Cycas), suggesting shared taxa among host populations within a region [15]; and no correlation was found between a resident cyanobiont species and its host cycad species Australia (*Macrozamia*) [16]. More recently a high diversity of endophytes in coralloid roots of the Asian cycad *Cycas bifida* was described using 16S rRNA Illumina amplicons [17]. Zheng et al. [17] found significant differences in the abundance of some bacterial families compared to regular cycad roots, with a dominance of cyanobacteria in the coralloid roots [17]. Likewise, Gutiérrez-García et al. [18] sampled the bacterial coralloid microbiome of *Dioon edule* using complete genomes and shotgun metagenomes from axenic cultures and liquid co-cultures that aim to describe the functional diversity of the endophytic microbiome [19], and also found several bacterial groups in addition to cyanobacteria endophytes. There appears to be a broad spectrum of non-cyanobacteria inside the coralloid root, which we are only beginning to explore.

Microbial communities associated with plants can be influenced by the plant compartment [20,21]. The soil surrounding plants can be considered as the main source of root endophytes [22]; the rhizosphere or soil region influenced by root exudates as a ‘growth chamber’ [22]; and the endosphere (internal tissues) as a restricted, specialized area that contains microbes [23]. In plants, comparison between soil and rhizosphere microbiomes often show differences in alpha [24] and beta diversity [24,25] and slight differences in taxonomic composition and community structure [26]. The endosphere typically has the lowest bacterial diversity [25], sometimes enriched with specific bacterial groups [27] and functionally specialized microbial communities [18]. In cycads, only two studies mention the influence of bacterial soil diversity in the bacterial coralloid endosphere. Zimmerman and Rosen, [10] found that endophytic *Nostoc* was different from edaphic *Nostoc* populations, while Cuddy et al. [28] found that the genotypes from cyanobacteria from the rhizosphere and the endophytes were different.

Our goal in this study is to describe the bacterial microbiome inside the coralloid root within the genus *Dioon*, as well as its surrounding soil and rhizosphere. We evaluated the bacterial diversity from seven *Dioon* species from a botanical garden population and a *Dioon merolae* from a natural population. Our main hypothesis was that there is bacterial diversity other than cyanobacteria in the coralloid roots microbiome, and that most of this diversity is recruited from the surrounding rhizosphere and enriched inside the root. This is the first study of the microbial diversity of a whole neotropical cycad genus including its rhizosphere and the surrounding bulk soil.

## Materials and Methods

### Field sampling and sample preparation

Coralloid roots samples of both juvenile (Fig 1) and adult plants (S1 Fig) were collected in April 2015 from the Botanical Garden “Francisco Javier Clavijero” from the Instituto Nacional de Ecología, A.C. in Xalapa (INECOL), Veracruz, Mexico (n=13); and from a natural population (n=4) from Santiago Lachiguiri, Oaxaca, Mexico. Bulk soil was collected (~40g) approximately 10 cm away from each plant from the botanical garden, although only three samples were fully processed (Table 1). Likewise, rhizosphere samples were collected from each plant from the botanical garden, of which only six were fully processed. All samples were transported to the laboratory in liquid nitrogen, frozen from the time of collection. The coralloid roots samples were defrosted and washed in a Phosphate Buffered Saline (PBS) for 20 minutes to obtain the rhizosphere for individuals from the botanical garden. Subsequently, to remove epiphytes, the coralloid roots were washed in hydrogen peroxide for seven minutes and immersed in 70% ethanol for 10 minutes. Then they were washed

**Fig 1.**
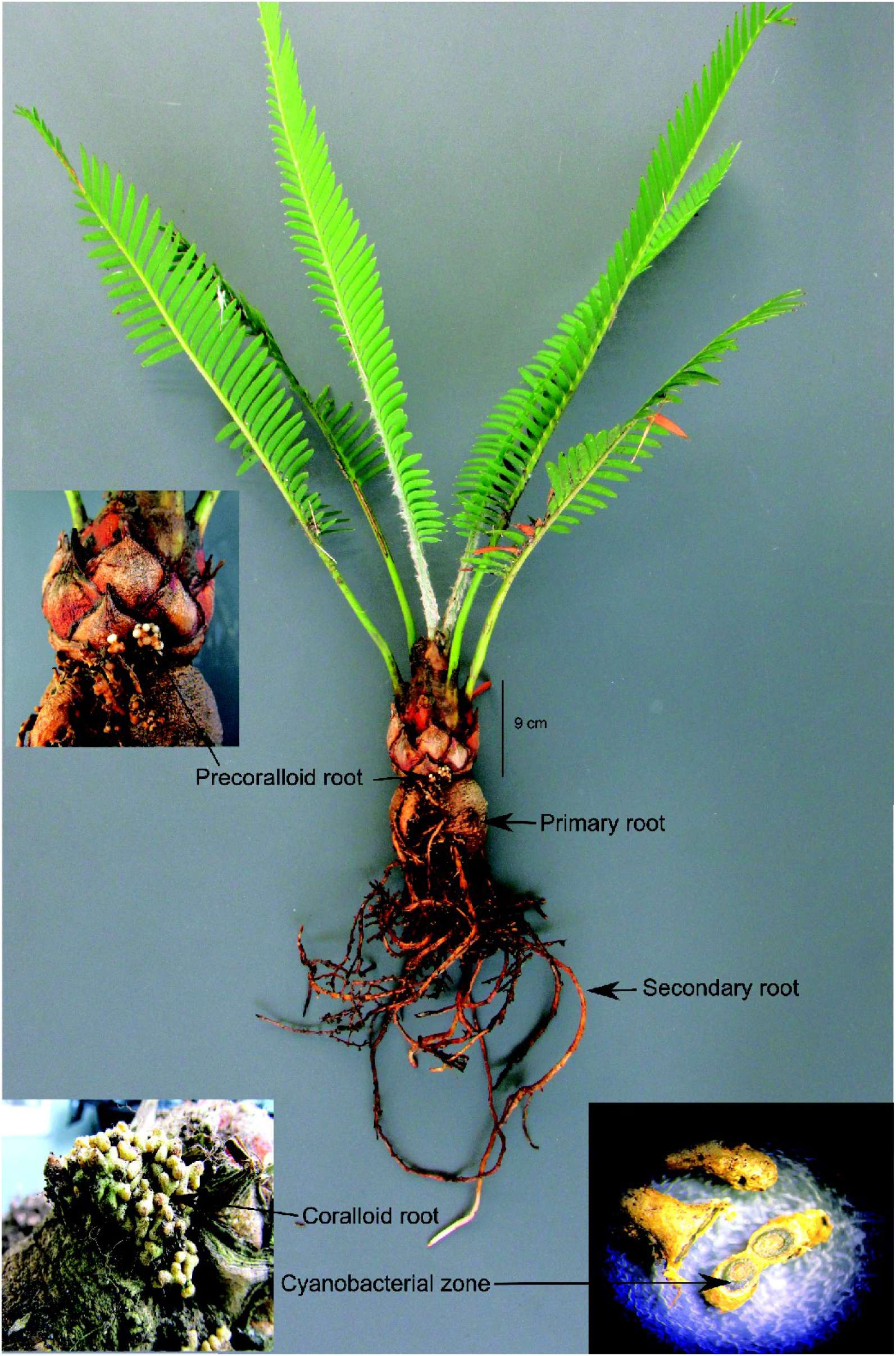
Juvenile individual of *Dioon edule* from the botanical garden. The radicular system of the *Dioon* species consist of a primary root, secondary root, precoralloid root and coralloid root. Also shown is a cross-section of the coralloid root without magnification, highlighting the ‘algal or cyanobacterial zone’.

**Table 1.** *Dioon* samples of different species, compartments, populations, and sets; The number of reads and OTUs obtained for each individual are also shown

| ID Sample | Species | Compartment | Population | #Set | #Reads | #OTUs |
| --- | --- | --- | --- | --- | --- | --- |
| 01.DME | *D. merolae* | END | BOG | 1 | 105344 | 1405 |
| 02.DME | *D. merolae* | END | BOG | 1 | 69583 | 1274 |
| 03.DME | *D. merolae* | END | BOG | 1 | 31357 | 1174 |
| 04.DME | *D. merolae* | END | BOG | 1 | 44118 | 2621 |
| 05.DME | *D. merolae* | END | BOG | 1 | 35857 | 1396 |
| 06.DME | *D. merolae* | END | BOG | 1 | 9424 | 482 |
| 01.DSO | *D. sonorensis* | END | BOG | 1 | 172972 | 4577 |
| 02.DSO | *D. sonorensis* | END | BOG | 1 | 11324 | 844 |
| 01.DAN | *D. angustifolium* | END | BOG | 1 | 28949 | 1303 |
| 01.DPU | *D. purpusii* | END | BOG | 1 | 60344 | 2177 |
| 01.DSP | *D. spinulosum* | END | BOG | 1 | 148326 | 1045 |
| 01.DED | *D. edule* | END | BOG | 1 | 6912 | 559 |
| 01.DMJ | *D. mejiae* | END | BOG | 1 | 3650 | 148 |
| 07.DME | *D. merolae* | END | NAP | 2 | 308911 | 417 |
| 08.DME | *D. merolae* | END | NAP | 2 | 339633 | 575 |
| 09.DME | *D. merolae* | END | NAP | 2 | 3293 | 583 |
| 10.DME | *D. merolae* | END | NAP | 2 | 56776 | 1187 |
| 01.DPU | *D. purpusii* | RHZ | BOG | 3 | 59691 | 3821 |
| 01.DSP | *D. spinulosum* | RHZ | BOG | 3 | 172698 | 4937 |
| 01.DED | *D. edule* | RHZ | BOG | 3 | 22247 | 1399 |
| 01.DME | *D. merolae* | RHZ | BOG | 3 | 13159 | 1692 |
| 01.DMJ | *D. mejiae* | RHZ | BOG | 3 | 93036 | 2837 |
| 06.DME | *D. merolae* | RHZ | BOG | 3 | 151769 | 4558 |
| 01.DPU | *D. purpusii* | BSO | BOG | 4 | 330178 | 6079 |
| 01.DSP | *D. spinulosum* | BSO | BOG | 4 | 52869 | 4139 |
| 01.DED | *D. edule* | BSO | BOG | 4 | 14903 | 2730 |

END = Endosphere; RHZ = Rhizosphere; BSO= Bulk soil, BOG= Botanical garden, NAP = Natural population

with 10% commercial bleach solution (NaClO) for 10 minutes, and then washed for five more times with sterile water before DNA extraction. Rinsed water from the last step in the washing process was tested for the presence of epiphytes by amplifying the 16S rRNA region, and also plated on axenic culture media (BHI medium). Only samples with no PCR amplification or culture growth were processed.

### DNA extraction and generation of 16S rRNA amplicons

Bacterial DNA was extracted from coralloid roots previously washed, using the DNeasy Plant mini kit (Qiagen). Approximately 120 mg of plant tissue from a sample was used for isolation of plant and bacterial DNA. The genomic DNA of rhizosphere and bulk soil samples were extracted using the UltraClean Soil DNA kit (MOBIO laboratories). We amplified the 16S rRNA hypervariable region V3-V4 using universal primers 515F and 806R [29] with 30 ng of gDNA. Only some soil and rhizosphere successfully amplified (Table 1). The amplicon libraries were prepared in our laboratory using the protocol by Vo & Jedlicka [30] without modifications and sequenced in a 2 x 300 bp paired-end run using the Illumina Miseq platform. The raw sequencing data for 16S rRNA sequencing results have been deposited at the Sequence Read Archive (SRA, http://www.ncbi.nlm.nih.gov/sra/) under accession numbers PRJNA481384 (https://www.ncbi.nlm.nih.gov/bioproject/PRJNA481384)

### Quality filters and sequence analysis

Forward and reverse sequences obtained from the MiSeq run were overlapped to form contiguous reads using PEAR v.0.9.8 [31] and were de-multiplexed using the QIIME v.1.9.1 pipeline [32] using a Phred score >Q30. Sequences were clustered into operational taxonomic units (OTUs) and taxonomy was assigned using UCLUST v.1.2.22q [33] based on 97% pairwise identity against the Greengenes database (gg_13_8_otus). Unassigned sequences were submitted to the Silva 16S rRNA database (release 132) in Mothur v.1.38.1 [34].

OTUs that belonged to the cyanobacteria phylum and that were initially assigned to this taxonomic level were blasted against another Silva database (release 123). OTUs assigned to chloroplast and mitochondria, and sequences that did not have at least ten counts across all samples were removed. Chimeric sequences, defined as hybrids between multiple DNA sequences, were identified using Mothur and also removed. The filtered OTUs were rarefied based on the number of sequences from the library with the lowest sequencing depth within each sets comparison, using QIIME.

### Taxonomic and phylogenetic diversity

We separated the 26 samples into sets (Table 1) to test the importance of the host species, the origin of the sample (natural vs botanical garden samples) and the root-associated compartment (endosphere, rhizosphere, and bulk soil). We first compared all *Dioon* species from the botanical garden endosphere samples (set 1, normalized count = 6912 reads), with the exception of a sample (*D. mejiae* 01.DJM) that had a very low number of reads (3650) and was removed. We estimated the alpha-diversity (observed species, Shannon and Simpson effective index) using the Rhea Pipeline v.2.0 [35] and the differences in the bacterial community within the sets (beta-diversity) were calculated with the Bray-Curtis distance [36]. Statistical tests for the alpha and beta diversity were not carried out in the set 1 comparison due the low number of samples for each species. To test for significant differences in alpha diversity we used a U-Mann-Whitney test for paired samples, and a Kruskal-Wallis test for comparisons of more than three samples. We also tested for significant differences in the beta-diversity of the communities using ANOSIM and PERMANOVA in R (VEGAN Package v1.17-2, [37]).

To measure taxonomic diversity among *Dioon* species from set 1, we used non-metric multidimensional scaling (nMDS) ordination with a Bray-Curtis distance matrix of taxon relative abundances, and used the balanced version of Unifrac distances, referred to as generalized UniFrac [38], to construct a dendrogram from all the samples using Ward’s clustering method [39]. We developed ‘heat trees’ with the taxonomic diversity found in the OTUs from set 1 (without 01.DMJ sample) and set 2 (normalized count = 3293 reads), defined as *D. merolae* samples from natural population, using the R package metacodeR v.0.2.1 [40]. In these plots, the node width and color indicate the number of reads assigned to each taxon. For each of these two sets, heat maps were performed for the 20 most abundant genera, using the function heatmap.2 from the R package gplots v.3.0.1 [41].

We compared only the *D. merolae* botanical garden samples from set 1 to their samples from a natural population (set 2) and tested for the differences in diversity among the endosphere and the rhizosphere (normalized count = 3650 reads), defined as set 3, from which where we expected most bacteria to be recruited. We also compared shared and private OTUs between all three compartments, soils samples, set 4, and their respective rhizosphere and endosphere samples (normalized count = 6912 reads), using alpha and beta diversity as described above. To visualize shared OTUs and taxa between the different comparisons we constructed Venn diagrams using R package VennDiagram v.1.6.20 [42]. In the set 4 comparison, we also wanted to know if endophytic cyanobacteria, a group known to be prevalent in coralloid roots, belonged to a single phylogenetic clade or they were members of different lineages. A single clade would suggest some specificity at the species or higher level, recruited from the rhizosphere and maintained or perhaps filtered, in the coralloid root endosphere. The cyanobacteria sequences that were classified to genus level and at least 400 bp in length from our study, along with a *Bacillus* sequence (outgroup) that was obtained from our study and reference sequences downloaded from GenBank database were aligned using Mafft v.7.305b [43]. The alignment was submitted to JModelTest v.2.1.10 [44] which determined that GTR + G +I model was the most appropriate substitution model. The phylogenetic estimation was performed using MrBayes v3.2.3 [45] with default parameters.

## Results

### Taxonomic diversity in the coralloid root endosphere

We analyzed the coralloid root-associated bacterial microbiome using 16S rRNA amplicons. Chimeras, chloroplast and mitochondria sequences and OTUs that did not have at least ten counts reads were discarded, resulting a total of 2,347,323 high-quality sequences and 8,812 OTU for the 26 samples (Table 1). The filtered OTUs were classified at the major taxonomic level possible and corresponded to 16 phyla, 40 classes, 78 orders, 137 families and 246 genera.

The taxonomic abundance profiles from the 12 *Dioon* samples from the botanical garden (set 1) were not grouped according to host species (Fig 2A), only 01.DME was more similar to 02.DME. Our dendrogram based on Ward’s clustering method with generalized UniFrac distance also showed a mixed distribution of hosts based on the total symbionts (Fig 2B). To measure the distribution of species diversity within the botanical garden, alpha diversity metrics (observed species/OTUs, effective Shannon and Simpson) were also estimated (S1 Table). Samples 01.DSO and 04.DME had the most alpha diversity, while 01.DSP and 06.DME were the least divers; In the latter two samples, *Nostoc* and Actinobacteria, dominate the community. Overall, these results suggest that in the botanical garden, all samples share a similar microbiome composition with varying degrees of diversity, independent of the host species.

**Fig 2.**
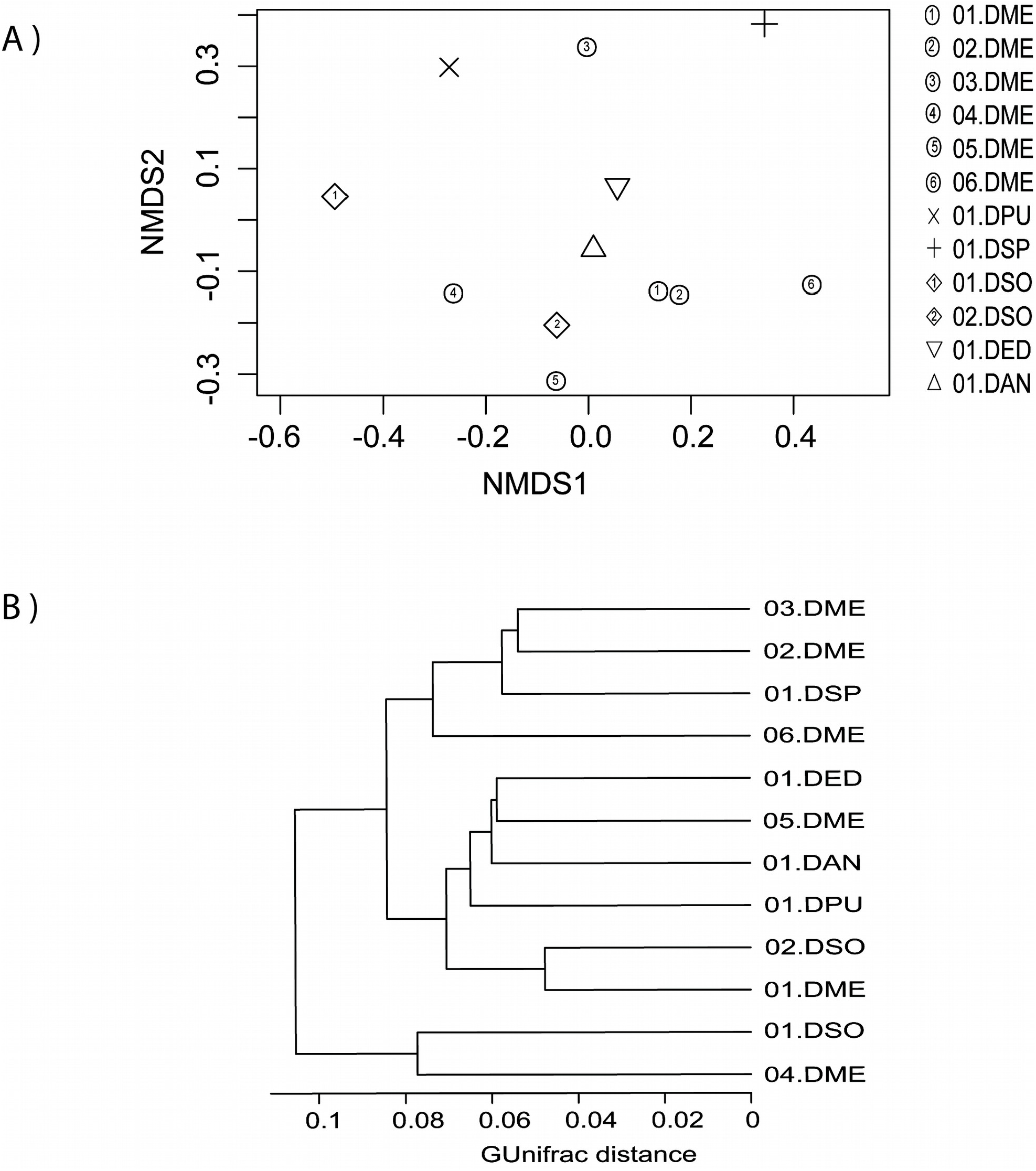
Beta-diversity analysis among the coralloid root endosphere from *Dioon* samples (set 1 comparison). A) Non-metric multidimensional scaling (nMDS) plot based on Bray-Curtis distance, where each symbol represents the bacterial community in a single *Dioon* sample. B) A dendrogram based on Ward’s clustering method showing the distance and clustering of the samples by their root endosphere microbiome. Abbreviations of the *Dioon* species used here are: DME = *D. merolae*, DPU= *D. purpusii*, DSP= *D. spinulosum*, DSO= *D. sonorense*, DED= *D. edule*, and DAN= *D. angustifolium*.

A sum of all the OTUs from the 12 botanic garden samples resulted in 4,156 OTUs, in which Proteobacteria, Actinobacteria, Cyanobacteria, Bacteroidetes, Verrucomicrobia were the five most abundant phyla (Fig 3A). The top five most abundant families were Nostocaceae, Streptomycetaceae, Rhizobiaceae, Pseudonocardiaceae, Chitinophagaceae. A total of 190 genera were identified in the *Dioon* endosphere (Fig 3A), and most of the 20 most abundant genera were evenly distributed among our samples, with a few notable exceptions (Fig 3B). *Nostoc, Rhizobium* and *Amycolaptosis* have distinctively high abundance for some samples (Fig 3B) (01.DSP, 05. DME and 06.DEM, respectively).

**Fig 3.**
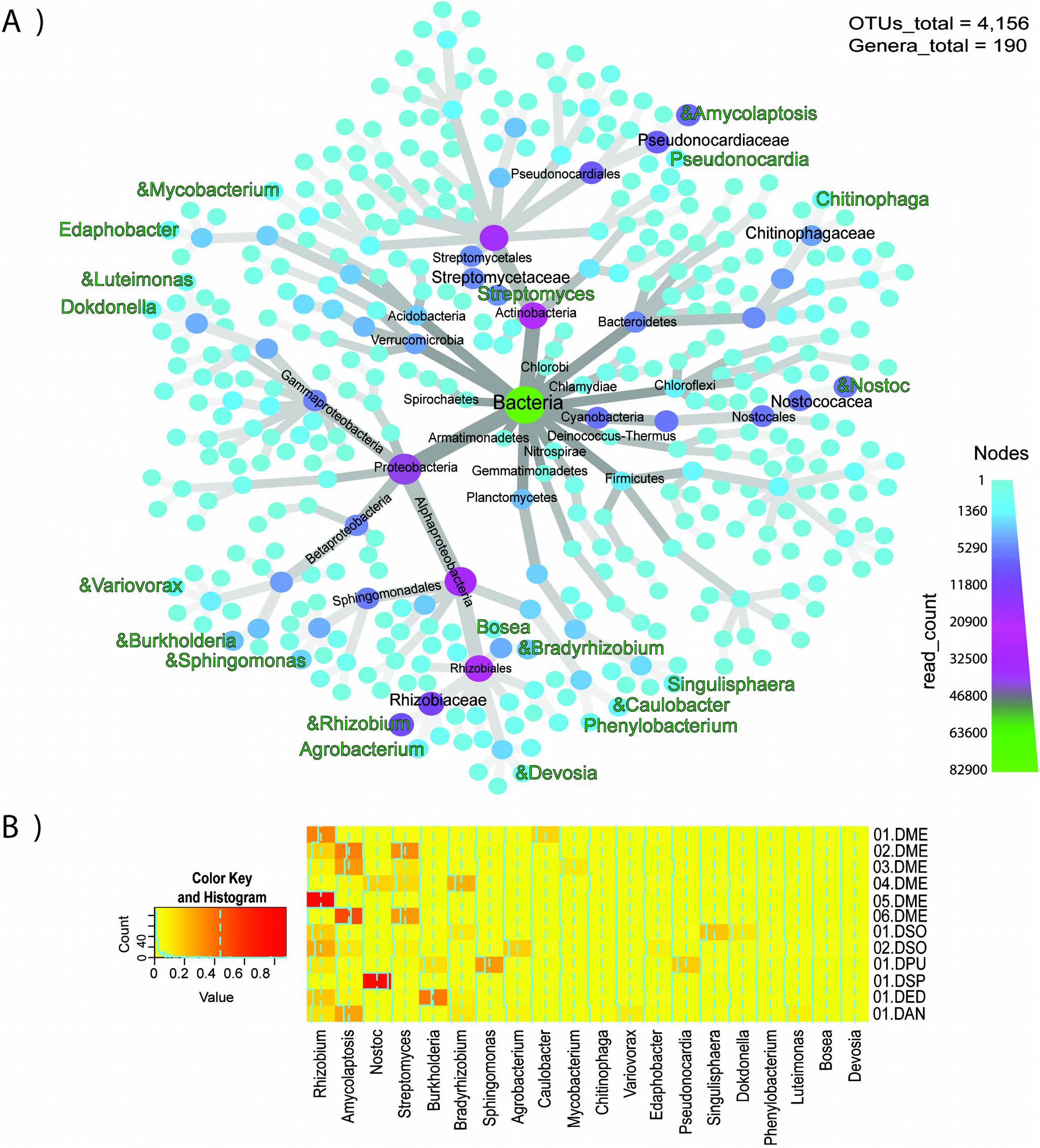
Taxonomic diversity and abundance of bacterial genera in the endosphere from *Dioon* samples (set 1 comparison). A) heat tree of the taxonomic diversity; The node width and color indicate the number of reads assigned to each taxon. Of the 20 most abundant genera (green) the symbol “&” represents the bacteria that have been reported as nitrogen-fixing. B) Heat map of 20 most abundant genera, where each column corresponds to a bacterial genus, and each row to a specific *Dioon* sample.

For *Dioon merolae* samples obtained from a natural population (set 2), there were a total of 902 OTUs and 91 genera (S2A Fig). The top five most abundant phyla were the same as the botanical garden samples except for Bacteriodetes. The five most abundant families were Nostocaceae, Streptomycetaceae, Geodermatophilaceae, Micromonosporaceae and Pseudonocardiaceae. The top most abundant genera included some that overlapped with samples from the botanic garden and were abundant in this latter population, namely *Nostoc, Rhizobium, Amycolatopsis, Streptomyces, Sphingomonas, Pseudonocardia, Mycobacterium, Bradyrhizobium, Agrobacterium* and *Devosia* (S2B Fig). Samples 07.DME and 08.DME from the natural population have less bacterial diversity and are mostly dominated by *Nostoc* (S2B Fig). The bacterial community of the natural population shared 445 (19%) OTUs and 77 (48%) genera with the botanic garden (S3 Fig). Differences of alpha diversity were not significant (p > 0.05, S2 Table), but beta diversity was (ANOSIM R= 0.7, p=0.004).

### Bulk soil and the rhizosphere as a source of the endophytes

We measured the shared and private bacterial diversity of the soil and compared it to the rhizosphere and the endosphere of three *Dioon* samples. There is high bacterial diversity in the three coralloid root compartments (5,740 OTUs and 207 genera), with soil being the most diverse compartment (3990 OTUs), followed by the rhizosphere and endosphere (3,411 and 1,537 OTUs, respectively) (Fig 4A). Differences in alpha diversity and beta-diversity were not significant in any of the comparisons (p > 0.05, S2 Table; p=0.18, PERMANOVA, respectively) with 635 OTUs and 84 genera shared among the three compartments. Yet the 20 most abundant shared genera are distributed differently in soil, rhizosphere or the endosphere (Fig 4B), with *Nostoc, Sphingomonas*, and *Rhizobium* among the most abundant genera. In the comparison of the three coralloid root compartments of the 20 most abundant shared genera, six genera were most abundant in the endosphere and five in the rhizosphere, and only the genus *Rhodobium* was most abundant in the bulk soil.

**Fig 4.**
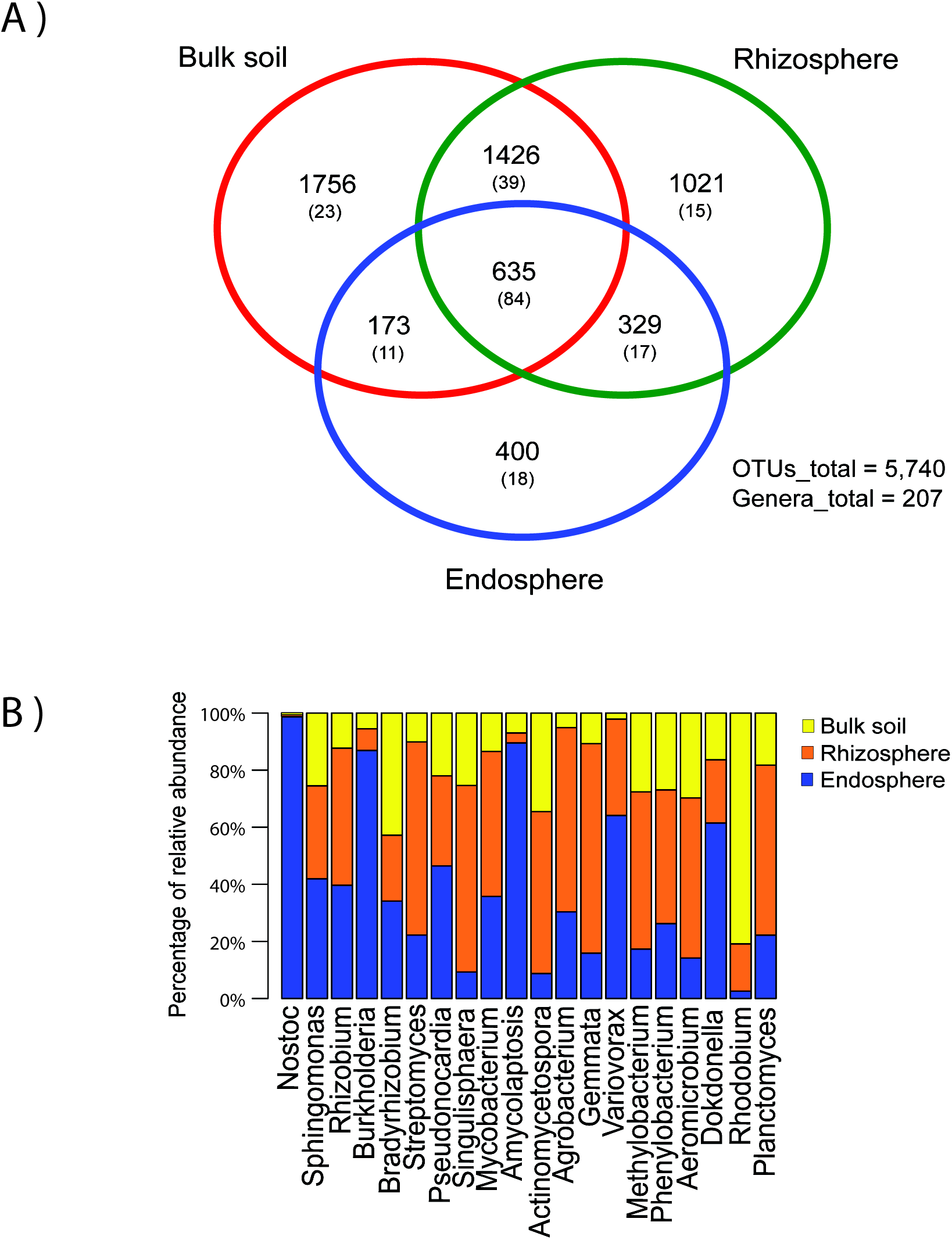
Bacterial diversity in the three coralloid root compartments (set 4 comparison). A) Venn diagram showing the shared OTUs and genera (in parenthesis) between the bulk soil, rhizosphere and endosphere. B) Bar chart of the 20 most abundant genera in the core of the three compartments (635 OTUs/84 genera).

Of the 1,537 OTUs and 130 genera found in the endosphere, 400 OTUs and 18 genera were private to this compartment, while 808 (53%) OTUs and 95 (73%) genera were shared with the bulk soil and 964 (63%) OTUs and 101 (78%) genera were shared with the rhizosphere. The outside (bulk soil and rhizosphere) and inside (endosphere) of the coralloid root shared 1137 (74%) OTUs and 112 (86%) genera. When comparing the rhizosphere of six *Dioon* samples (set 3) with their respective endosphere, we found 4219 OTUs and 187 genera, of which 1044 (65%) OTUs and 116 (62%) genera were shared (Fig 5A), while 574 OTUs and 23 genera were private to the endosphere. There is significantly more bacterial diversity in the rhizosphere for the three metrics of alpha diversity (Fig 5B, S2 Table); although beta-diversity was not significantly different (p=0.09, PERMANOVA).

**Fig 5.**
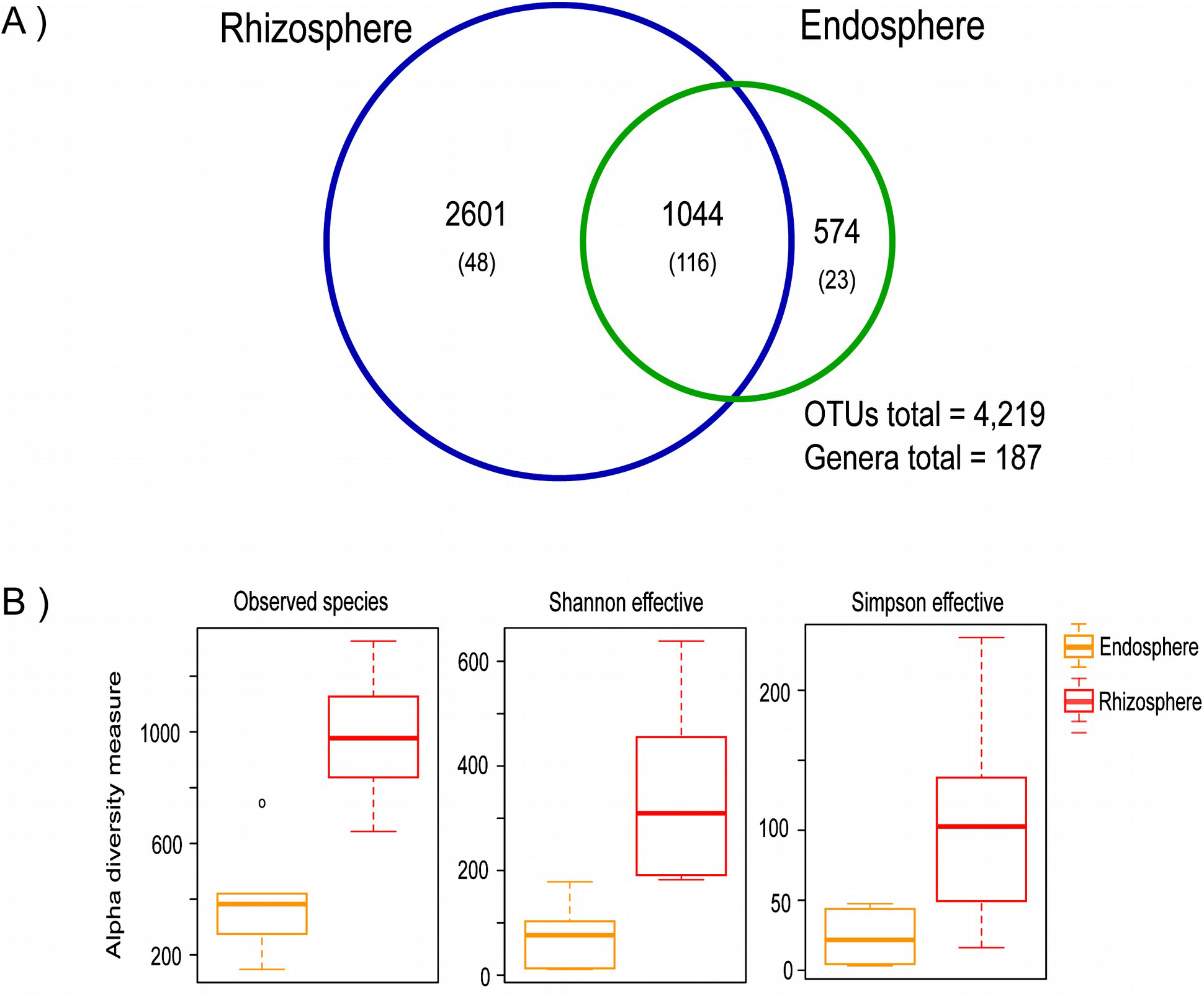
Differences of bacterial diversity between the endosphere and rhizosphere (set 3 comparison). A) Venn diagram showing the shared OTUs and genera (in parenthesis) between the endosphere and rhizosphere of the six *Dioon* samples. B) Distribution of alpha-diversity within the rhizosphere and endosphere as measured by observed species, Shannon and Simpson effective index.

### Phylogeny of the cyanobacterial endophytes

When we compared the cyanobacteria microbiome from the three compartments associated to coralloid root, we found 64 OTUs for the phylum, of which 38 can be classified to seven families and in seven genera: *Nostoc, Calothrix, Microcoleus, Leptolyngbya, Chroococcus, Acaryochloris, Scytonema*. To test for possible phylogenetic patterns that would suggest specificity to the coralloid root compartment, we placed our samples in a phylogeny constructed using 25 OTUs classified to the genus level (400 bp) (Fig 6).

**Fig 6.**
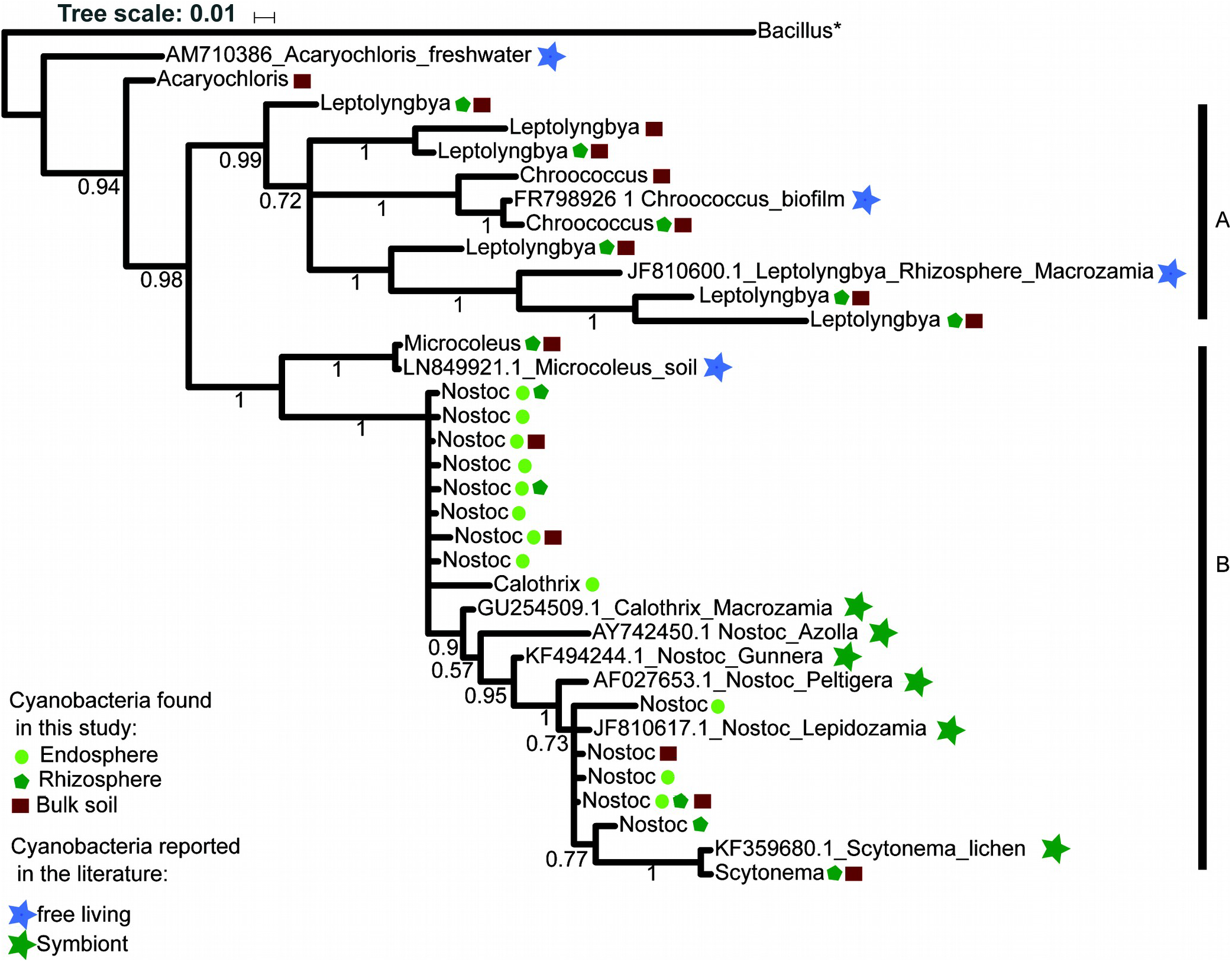
Phylogeny of the genera belong to cyanobacteria found in the three compartments (set 4 comparison). The ingroup contains 7 cyanobacteria genera and 25 OTUs. The endosphere, rhizosphere, and bulk soil are represented by a circle (yellow), Pentagon (Maroon), and square (Green) respectively. The cyanobacteria reported in the literature are represented by a star, the color blue indicates free-living cyanobacteria and green indicates those that live in symbiosis. The letters: A and B denote the main clades formed in the analysis. The values above branches are the posterior probabilities.

The resulting tree shows two clades, with the genus *Acaryochloris* as the sister taxon to a clade of all other genera obtained from our amplicons data. Clade (A), included two OTUs classified as the genus *Chroococcus* and six OTUs as *Leptolyngbya* found in the bulk soil and/or the *Dioon* rhizosphere. They grouped with a previously reported *Chroococcus* obtained from a biofilm of a water fountain; and most notably, a *Leptolyngbya* isolated from a distantly-related cycad’s rhizosphere (*Macrozamia sp*. ‘Bundarra’ from Australia, Cuddy et al. [28]). Clade (B) included a subclade with only one rhizosphere and soil OTU identified as *Microcoleus;* and a subclade with samples from both cycad endophytes and the cycad rhizosphere and bulk soil classified as *Nostoc, Calothrix*, and *Scytonema*. Some of these genera include previously reported endosymbiotic *Nostoc* in cycads and early seed plants (*Gunnera*) and the aquatic fern (*Azolla*).

Interestingly, all of our Nostocales cycad endophytes constitute a monophyletic group that likely include a new species of *Nostoc* or perhaps even a new genus. It is worth mentioning that *Nostoc* and *Calothrix* genera were the only Nostocales genera found in the endosphere among the 2,347,323 reads and 8,812 OTUs from the 26 individuals included in this study. *Calothrix* was exclusive to the endosphere.

## Discussion

### A taxonomically diverse bacterial community inside *Dioon* coralloid roots

Endophytes associated with cycad coralloid roots have been studied for more than a century mostly by methods that underestimated or did not aim to measure the non-cyanobacterial bacterial diversity present inside the root. Our study supports very early studies [4,8,9], and two more recent studies, the 16S rRNA-based Zheng et al. [17] and the metagenomic-based Gutiérrez-García et al. [18], that report a diverse bacterial microbiome beyond cyanobacteria.

We found a high bacterial diversity in the endophytes of the 12 samples from six Mexican *Dioon* species, similar to that detected in the root endosphere from angiosperm plants as *Arabidopsis* [27], sugarcane [46], poplar trees [47], and inside the specialized root nodules of legumes [48]. We found that Proteobacteria, Actinobacteria, Cyanobacteria, Bacteroidetes and Planctomycetes were the dominant bacterial phyla in the *Dioon* coralloid root microbiome. Some of these phyla (Proteobacteria, Actinobacteria and Bacteroidetes) have been reported as typical root endophytes, enriched in other plants [49].

At the level of family Nostocaceae is among the most abundant as also reported in *Cycas bifida* [17]. We also found the 14 other most abundant families from *C. bifida* that are present in our *Dioon* samples. In the Gutiérrez-García et al. [18] study, the authors used shotgun metagenomics to characterize co-cultures from an inoculum of the coralloid root endosphere from *D. merolae* and found 51 families and 76 bacterial genera, of which 34 and 33 respectively, were present in our amplicon data. Nine families were shared between these two studies and ours, including Nostocaceae, Rhizobiaceae, Pseudonocardiaceae, Mycobacteriaceae, Hyphomicrobiaceae, Bacillaceae, Bradyrhizobiaceae that were abundant in our data, suggesting these may constitute a taxonomic core for cycads in general.

Most of the diversity was evenly distributed among most samples, with the exception of three samples which showed that cyanobacteria were highly abundant and dominant over other taxa. The overall low bacterial diversity associated with a high cyanobacteria abundance was also observed in three coralloid roots samples from *Cycas bifida* [17]. We partially agree with the authors that cyanobacteria may inhibit growth of other groups by competition or secretion of secondary metabolites. We also suggest however, that it is possible that the age of the coralloid root is an important factor influencing the dominance of cyanobacteria, and that older roots could contain more cyanobacteria as they mature. The maturation of a coralloid root causes morphological changes that include more numerous lenticels [3] and could therefore allow the entry of more bacteria. Cyanobacteria colonization is important in the transition from precoralloid to coralloid root [3], and in addition, changes in the structure of the cyanobacterial community have been reported during the degradation of the coralloid roots [50]. The age of the root as a key factor for microbiome composition inside the coralloid root requires further investigation.

### Composition and bacterial diversity of the root-endosphere microbiome among sites and host species

Differences in the growth stage of the natural (adults) and botanic garden (juveniles) host samples could be affecting the beta-diversity in the root microbiome, as is seen in other plants such as legumes [48] and rice [51]. Physiological and biological differences among the hosts could also affect the composition and quantity of root exudates throughout the plant growth stages [52,53], influencing the diversity of the root endophytes and the microbial conformation of the rhizosphere [54]. The varying microenvironmental factors in each population can also influence in the prokaryotic communities from the two sites that we compared. Biogeographic factors significantly affect the microbial communities associated in roots from *Agave* [20] and the soil type and geographical location contribute to rice root microbiome variation [25]. Future samples from natural populations including their rhizosphere will help discern the importance of local climatic and soil conditions in the endophyte taxonomic diversity.

For most samples, there is no clear relationship to the host species. This has been observed in other plants; for instance, there were no differences in the alpha and beta diversity of microbial communities from roots of three species of *Fragaria* strawberries [55]. The sample tissue type (different plant compartments) accounted for more of the variance in the prokaryotic communities than the host species in cultivated and native *Agave* species [20]. Other variables such as soil type have been shown to have more influence than host genotype [25] or plant species [56] on the endosphere microbiome. Furthermore, similar genus-level microbiomes have been observed for different genera from legumes [48] and maize and teosinte in Poaceae [57]. Finally, it is also possible that given that our samples seem to be part of a species complex of recent divergence [58,59], they simply share biological traits that favor certain bacterial communities.

Our data also show that there are only a few bacterial genera enriched in the *Dioon* endosphere. More than half of the most abundant genera we found have been reported as diazotrophic plant endophytes, confirming the nitrogen-fixation role of the coralloid root [60]. Interestingly, the Rhizobiales were the most abundant order in the coralloid root microbiome. Rhizobiales have been reported in nodules [48,61] and roots [61] of legumes, and roots of non-legume plant species as sugarcane [46] and *Agave* [62]. The presence and enrichment of Rhizobiales species in the roots of phylogenetically diverse plant hosts [63] could indicate that the Rhizobiales are part of a bacterial core in symbiotic plant microbiomes. Diverse species of the rhizobiales can affect the primary root growth in non-legume plants as *A. thaliana* [63] and are a key component in the formation of nodules in different species of legumes [64]. It remains to be seen if their function in the coralloid root is similar to their functions in angiosperms.

Some genera among the 20 most abundant such as *Agrobacterium, Bosea, Bradyrhizobium, Burkholderia, Caulobacter, Chitinophaga, Devosia, Luteimonas, Mycobacterium, Rhizobium, Sphingomonas, Streptomyces, Variovorax* are also known to have traits of plant growth-promotion [65–67].Plant growth-promoting bacteria were also consistently enriched in the two *D. merolae* populations, and include *Streptomyces, Sphingomonas, Rhizobium, Pseudonocardia, Nostoc, Mycobacterium, Bradyrhizobium, Amycolatopsis*, and *Agrobacterium*. Samad et al. [66] isolated the three most abundant genera found in their amplicon data from four weed and grapevine root microbiomes and found that *Sphingomonas* have beneficial properties such as auxin (indole acetic acid) production. It is thus possible that the plant is favoring some beneficial bacterial groups as endophytes. In summary, the prevalence of these bacterial groups suggests a functional core that could be important in the development and growth of the coralloid root.

### The bulk soil and rhizosphere serve as a bacterial reservoir for the colonization of endosphere

Our results demonstrated that the soil and rhizosphere associated with *Dioon* coralloid roots are highly diverse. As with other plants [25,66,67], the soil (bulk soil and rhizosphere) is more diverse in the number of OTUs and genera than the root endosphere, although these differences are not significant in our study (alpha diversity metrics). When we compared the rhizosphere and endosphere from the six *Dioon* samples (set 3 comparison), we found significant differences in the alpha diversity, with higher diversity on the outside than the inside of coralloid root, a common pattern reported previously in 30 species of angiosperms [68].

The lack of significant differences in beta diversity among the samples (set 3 and set 4) with their respective endospheres support the idea that the soil microbiota is the main source of species for the root endosphere. This is similar to what has been reported for sugarcane, in which the 90% of OTUs discovered in the endosphere of roots, stalks, and leaves were also present in the bulk soil samples [46]. In *Agave* species leaf and root bacterial endophytes were present in their respective episphere (93%) and bulk soil (86%) [20]. Finally, with respect to the taxa from the endosphere that are not shared with the rhizosphere and bulk soil, only the genus *Bacteroides* was not found in the soil, reported previously as an endophyte of rice (*Oryza*) [69] and reed (*Phragmites australis*) [70].

The colonization of bacteria from the rhizosphere to the endosphere may take place through the cracks or lenticels in coralloid roots as an active process [3,4,71], as many endophytes express cell-wall-degrading enzymes [72]. Further research could investigate the function (metagenomic) and gene expression patterns (metatranscriptomic) associated with the coralloid root colonization by the cycad endophytes we report here.

### The endosphere of *Dioon* is enriched with cyanobacteria within a monophyletic group

Our study finds various taxa that belong to the cyanobacteria phylum in the *Dioon* coralloid root-associated microbiome. These include 38 OTUs grouped in seven genera, of which four (*Microcoleus, Chroococcus, Acaryochloris, Scytonema*) had been not been previously reported in soils surrounding the coralloid root.

Our phylogeny shows that *Dioon* Nostocales endophytes are close relatives, placed in the cyanobacteria heterocystous cluster [73]. Genera such as *Scytonema* are found in the rhizosphere and bulk soil and we do not rule out the possibility that other non-heterocystous genera capable of nitrogen fixation such as *Leptolyngya* [74] could colonize the endosphere. Our results, however, show that *Nostoc* and *Calothrix* are the predominant two genera inside the coralloid root. They have heterocysts, thick-walled cells where the nitrogenase enzyme for nitrogen fixation is located [75] and are known for their colonization ability and for promoting the growth of several plant groups [76,77]. Earlier studies of the cycad coralloid root endophytes also mostly report a bacterial community composed of a few or many strains of *Nostoc*, and less frequently, *Calothrix* [9,11,14]. The genera *Tolypothrix* and *Leptolyngbya* found in the soil surrounding our *Dioon* samples, were also previously reported in the roots from the Australian *Macrozamia* [28] *Tolypothrix* was not found in our sequences, perhaps due to inconsistencies in taxonomic reports in this genus [78].

It remains to be seen if the prevalence of only some bacterial groups in the endosphere is a result of plant-driven selection mechanisms through the enhanced activity of defense and hydrolytic enzymes in the plant host [77], or it is due to the coralloid root microenvironment such as the influence of soil pH and nitrates content in the bacterial diversity associate to roots of soybean and wheat [67]; or the microbiome interactions themselves that could filter out certain bacteria groups including cyanobacteria [22,79].

Also, the finding of similar species in the rhizosphere opens the possibility that species assemblages that already exist outside the root, enter as such when recruited. The finding of interacting *Nostoc-Caulobacter* in other *Dioon* species adds to this possibility [18]. These hypotheses require appraisal based on a wider sampling of Mexican cycad species and functional studies of the main groups we identified here.

## ACKNOWLEDGMENTS

We would like to thank F. Pérez-Zavala, P. Pelaez-Hernández, and M. Pérez-Farrera for invaluable assistance during field work. We also thank J.C. García-Sánchez for his help in the preparation up of the amplicon libraries.

## Supporting information

**S1 Table. Alpha diversity metrics for the 12 samples of genus *Dioon* (set 1).**

**S2 Table. Alpha diversity pairwise comparisons among sets of populations and compartments.**

**S1 Fig. Adult individual of *Dioon merolae* from the natural population.**

**S2 Fig. Taxonomic diversity and abundance of bacterial genera in the endosphere from *Dioon merolae* (set 2).** A) Heat tree of taxonomic diversity, the node width and color indicates the number of reads assigned to each taxon. Of the 20 most abundant genera (green) the symbol “&” represents the bacteria that have been reported as nitrogen-fixing. B) Heat map of 20 most abundant genera, each column corresponds to a bacterial genus, each row to a specific *Dioon merolae* sample.

S2 Fig. Venn diagram showing the shared OTUs and genera (in parenthesis) between the *D. merolae* samples from botanical garden and natural population.

**S3 Fig. Differences of bacterial diversity between the two populations from *D. merolae* (set 2 comparison).** Venn diagram showing the shared OTUs and genera (in parenthesis) between the *D. merolae* samples from botanical garden and natural population.

